## Supplementary Figures/Tables/Movie legends for "Autonomous and non-cell autonomous etiology of ciliopathy associated structural birth defects"

### Table S1. *Midgestation lethality of Ift140^null1/null1^ embryo*

**+/+	+/-	-/-	-/-***

E9	15	30	23	0

E10	18	16	20	0

E11	7	20	10	2

E12	16	26	15	5

E13	19	54	15	10

E14	17	30	5	2

E15	9	14	2	1

E16	5	2	0	0

*Subset of -/- embryos that died (no heartbeat or blood flow, or already undergoing resorption.

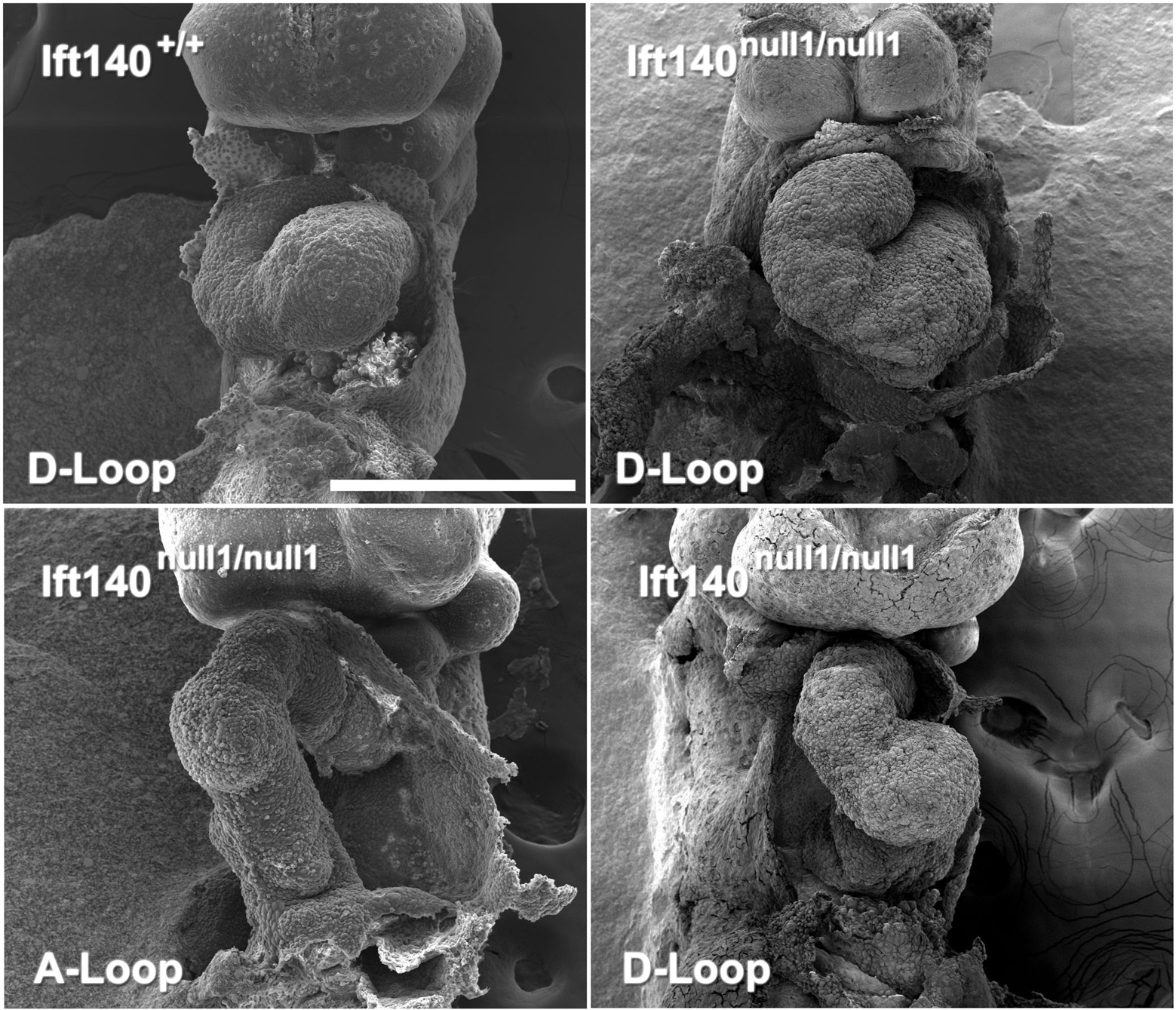

**Figure S1. Scanning EM of E9 hearts.** Scanning electron microscopy of a litter of *Ift140^null1^* embryos. Scale bar is 0.5 mm.

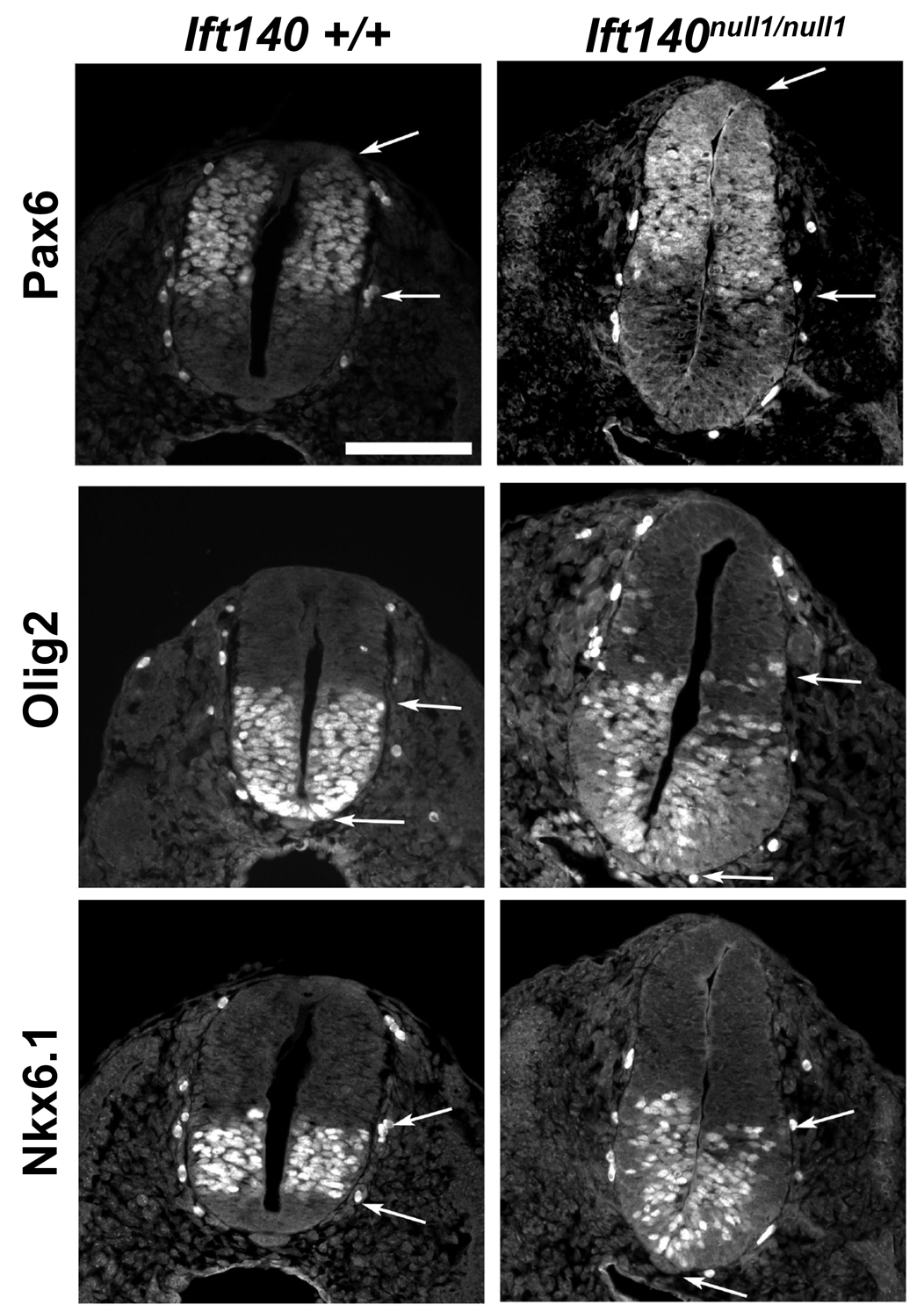

**Figure S2. Immunostaining for differentiation markers Olig2, Nkx6.1, and Pax6, in the neural tube of E10.5 in the Ift140^null1/null1^ embryos**.

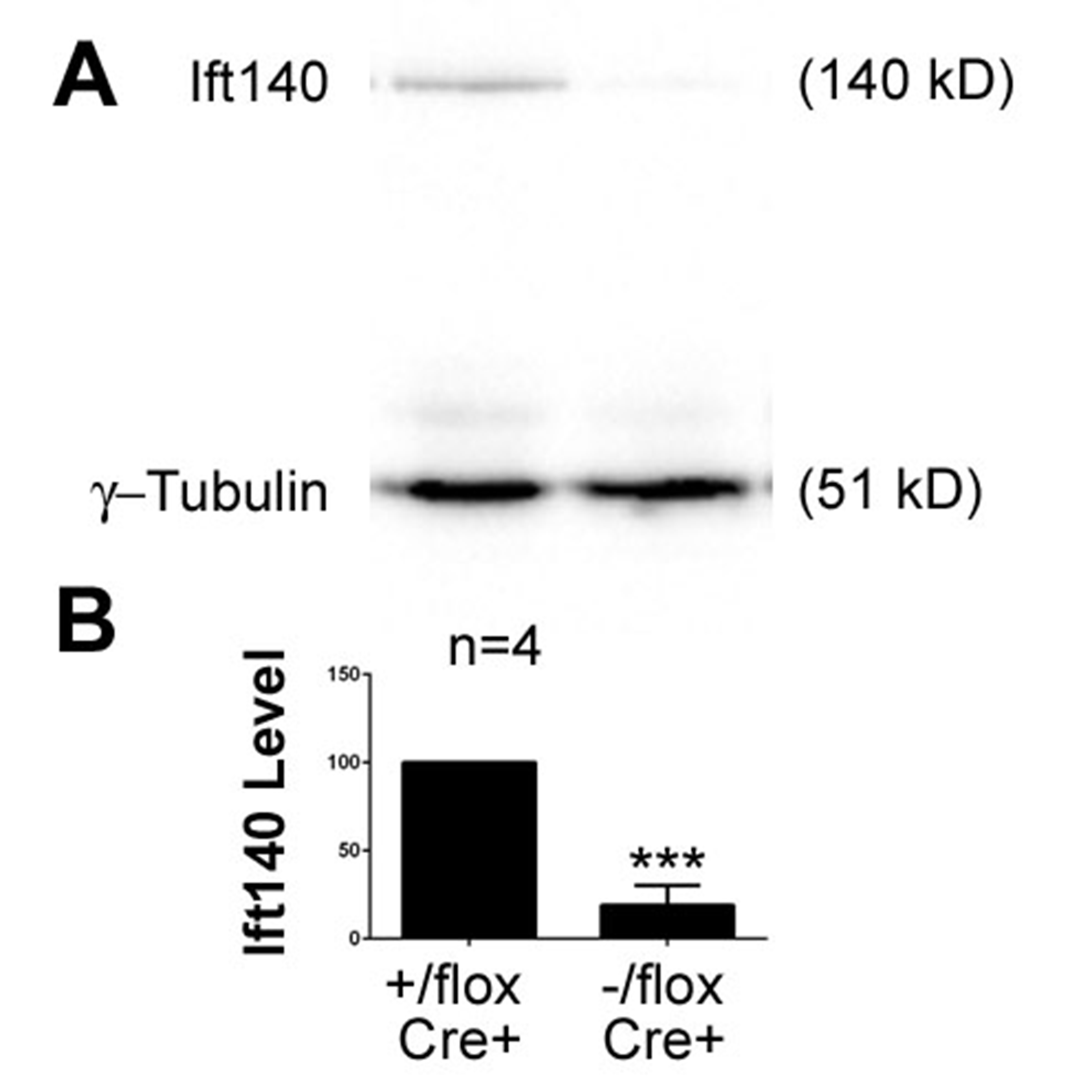

**Figure S3. Loss of Ift140 after tamoxifen treatment.**

**A.** Western blot showing the level of Ift140 in embryos 48 hr after treatment of the mother with tamoxifen. γ-tubulin is a loading control.

**B.** Quantification of the extent of Ift140 reduction 48 hr after treatment of the mother with tamoxifen. Embryos were treated at E9 and harvested at E11. Level of Ift140 was normalized between embryos using γ-tubulin and then experimental and control embryos within a litter were ratioed with controls set to 100%. N=4 experimentals and 4 controls. ***p<0.0001. Error bar is standard deviation.

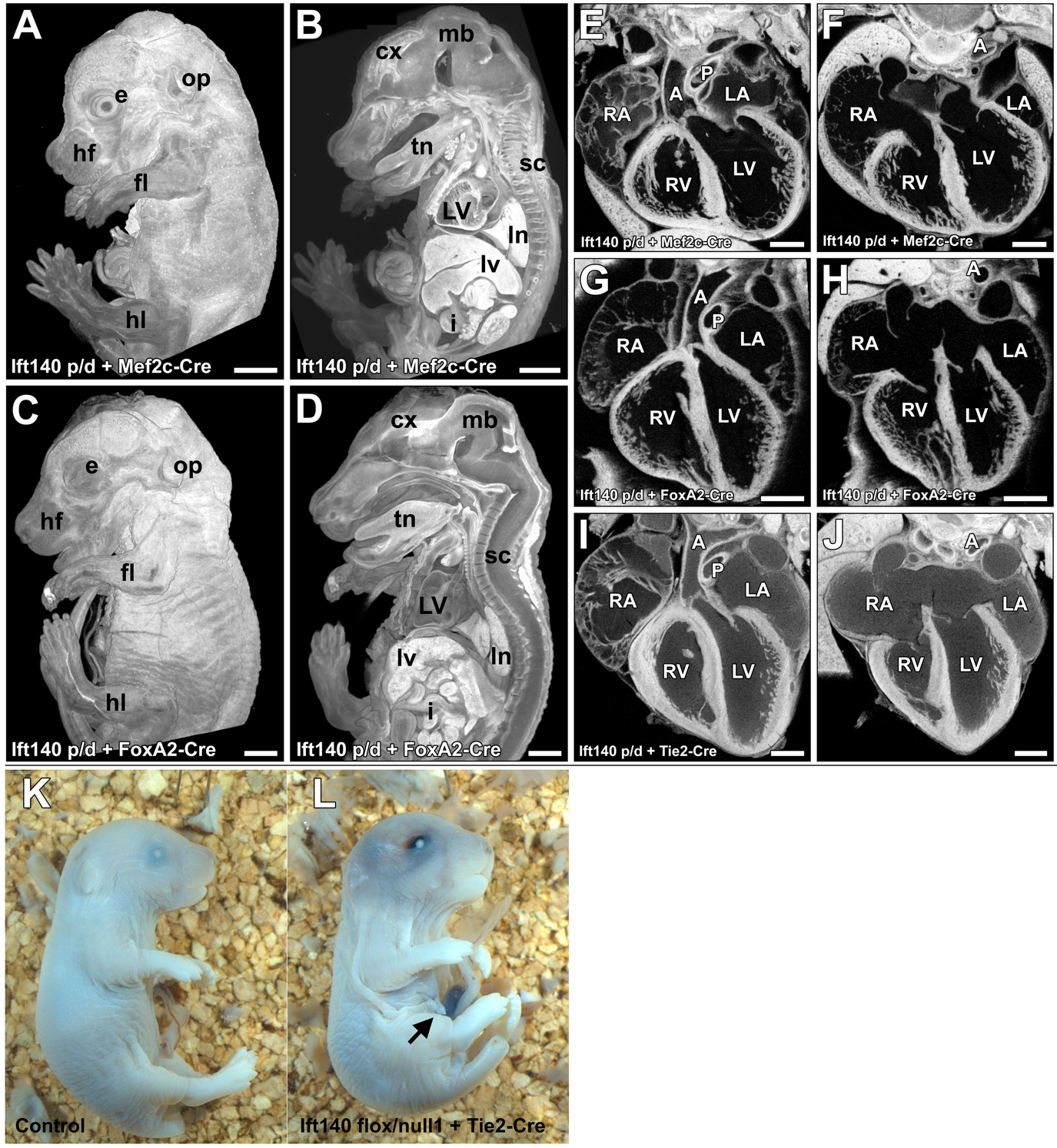

**Figure S4. Mef2c-Cre, Tie2-Cre, or tamoxifen-driven FoxA2-CreER deletion of Ift140 does not cause extensive cardiac phenotypes. A-D**. Deletion of *Ift140* by Mef2c-Cre or by tamoxifen induced FoxA2-CreER (tamoxifen administered at E6.5, E7.5 or E8.5) display normal whole body gross anatomy. **E-J**. Deletion of *Ift140* by Mef2c-Cre, Tie2-Cre, or by tamoxifen driven FoxA2-CreER did not affect cardiac and great vessel anatomy. **K, L.** Deletion of *Ift140* by Tie2-Cre results in eyelid closure defects and supernumerary mammary glands (arrow). LV: left ventricle; cx: cerebral cortex; sc: spinal cord; mb: midbrain; fl: forelimb; hl: hindlimb; e: eye; op: otic placode; hf: hair follicles; lv: liver; ln: lungs; i: small intestine; A: aorta; P: pulmonary trunk; LV: left ventricle; RV: right ventricle; LA: left atria; RA: right atria. Scales bars: **A-D** = 1 mm, **E-J** = 0. 5 mm.

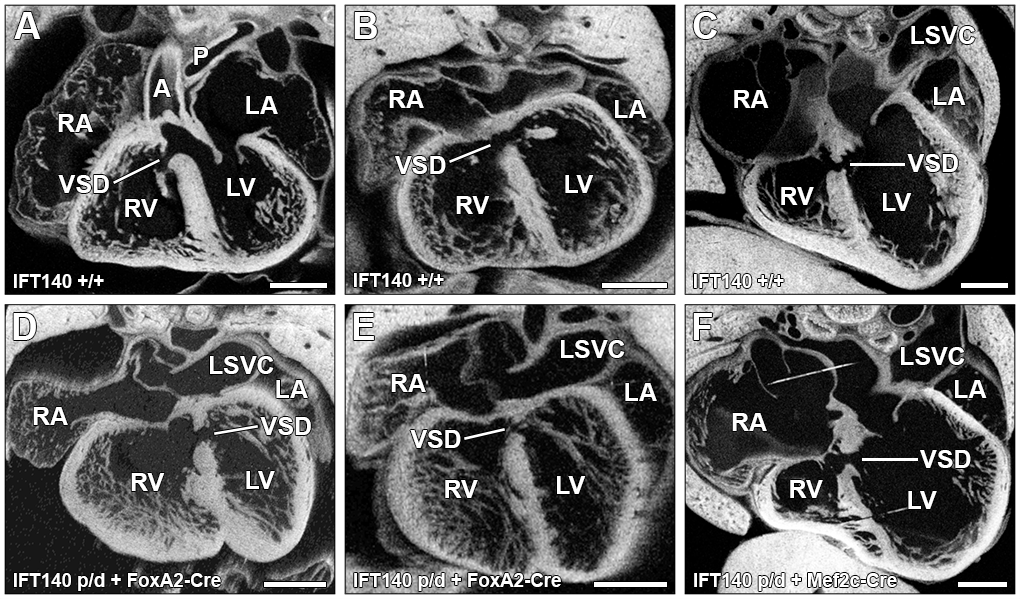

**Figure S5. Tamoxifen treatment appears to cause small ventricular septal defects. A-C.** A small but significant number of littermate controls collected from tamoxifen treated litters were found to have small ventricular septal defects (VSD). **D-F.** Similar small ventricular septal defects were also seen in a small number of embryos with tamoxifen driven Cre specific knockdown, including FoxA2-Cre (D,E) and Mef2c-Cre (F). As these defects were seen across both wild type and experimental knockdown groups they were excluded from phenotypic analysis and categorized as experimental artifacts. A: aorta; P: pulmonary trunk; LV: left ventricle; RV: right ventricle; LA: left atria; RA: right atria; VSD: ventricular septal defect; LSVC: left superior vena cava.

All scales bars = 0.5 mm.

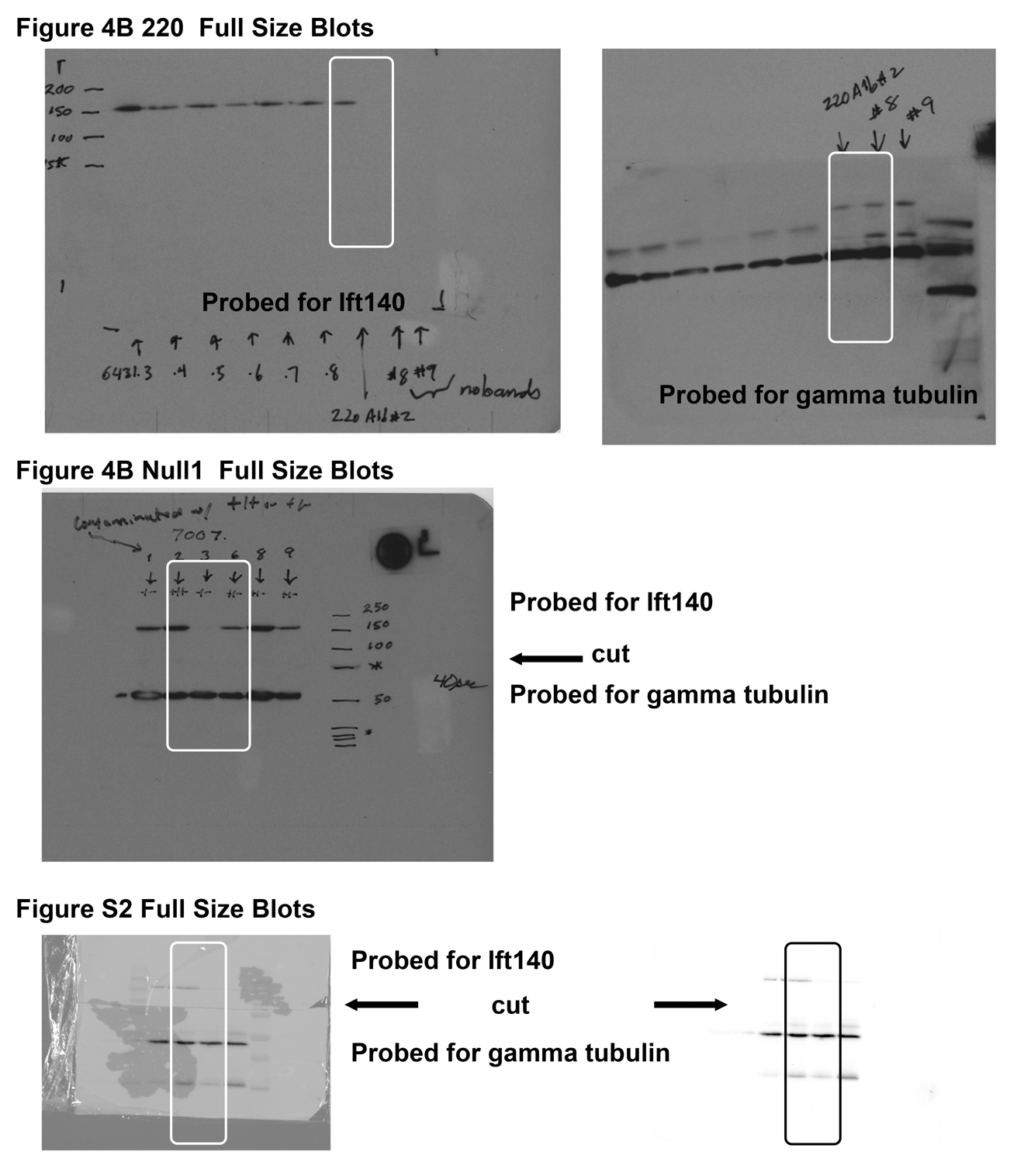

**Figure S6. Full Size Western Blots. A.** Left membrane was probed for Ift140 while the right was probed for gamma tubulin. The lanes used in Figure 4B are in the white box. **B.** Membrane was cut and the top half probed for Ift140 and the lower half probed for gamma tubulin. The lanes used in Figure 4B are in the white box. **C.** Membrane was cut and the top half probed for Ift140 and the lower half probed for gamma tubulin. The left image is the western blot superimposed on an image of the membrane while the right image is only the western blot. The lanes used in Figure S2 are in the boxes.

**Supplementary Movie 1.**

3D reconstruction of a wildtype E16.5 embryo processed using episcopic confocal microscopy highlighting normal cardiac anatomy. Ao: Aorta, dAo: Descending aorta, DA: Ductus arteriosus, LA: Left atria, LV: Left ventricle, MV: Mitral valve, PA: Pulmonary artery, PT: Pulmonary trunk, RA: Right atria, RV Right ventricle, TV: Tricuspid valve.

**Supplementary Movie 2.**

3D reconstruction of a E16.5 IFT140^null1/null1^ embryo processed using episcopic confocal microscopy highlighting abnormal cardiac anatomy. AVSD: Atrioventricular septal defect, CA: Common atria, L: Liver, LA: Left atria, LV: Left ventricle, PTA: Persistent truncus arteriosus, RA: Right atria, RV Right ventricle, SC: Spinal cord.

**Supplementary Movie 3.**

3D reconstruction of a wildtype E16.5 embryo processed using episcopic confocal microscopy highlighting normal cardiac outflow tact development. Ao: Aorta, dAo: Descending aorta, ASLV: Aortic semilunar valve, DA: Ductus arteriosus, LA: Left atria, LB: Left Bronchus, LPA: Left pulmonary artery, LV: Left ventricle, PA: Pulmonary artery, PSLV: Pulmonary semilunar valve, PT: Pulmonary trunk, RA: Right atria, RB: Right Bronchus, RCA: Right carotid artery, RPA: Right pulmonary artery, RV Right ventricle, T: Trachea.

**Supplementary Movie 4.**

3D reconstruction of a E16.5 IFT140^null1/null1^ embryo processed using episcopic confocal microscopy highlighting abnormal cardiac outflow tact development. dAo: Descending aorta, AVSD: Atrioventricular septal defect, CA: Common atria, E: Esophagus, IVC: Inferior vena cava, L: Liver, PTA: Persistent truncus arteriosus, RV Right ventricle, S: Stomach, TEF: Tracheoesophageal fistula, UL: Underdeveloped Lung.

**Supplementary Movie 5.**

3D reconstruction of a wildtype E16.5 embryo processed using episcopic confocal microscopy highlighting normal Trachea/Esophagus development. Ao: Aorta, dAo: Descending aorta, E: Esophagus, LA: Left atria, LB: Left Bronchus, LCA: Left carotid artery, LV: Left ventricle, MV: Mitral valve, PV: Pulmonary vein, RA: Right atria, RB: Right Bronchus, RCA: Right carotid artery, RV Right ventricle, S: Stomach, SCV: Subclavian vein, T: Trachea, TV: Tricuspid valve, VC: Vena cava.

**Supplementary Movie 6.**

3D reconstruction of a E16.5 IFT140^null1/null1^ embryo processed using episcopic confocal microscopy highlighting Tracheoesophageal fistula. E: Esophagus, IVC: Inferior vena cava, L: Liver, S: Stomach, SC: Spinal cord, SCV: Subclavian vein, SVC: Superior vena cava, TEF: Tracheoesophageal fistula, UL: Underdeveloped Lung.

**Supplementary Movie 7.**

3D reconstruction of a wildtype E16.5 embryo processed using episcopic confocal microscopy highlighting normal chest/lung development. dAo: Descending aorta, D: Diaphragm, E: Esophagus, L: Liver, LB: Left Bronchus, LL: Left lung lobe, RB: Right Bronchus, RL(SL): Right lung (Superior lobe), RL(ML): Right lung (Middle lobe), RL(IL): Right lung (Inferior lobe), RL(PCL): Right lung (Post-caval lobe).

**Supplementary Movie 8.**

3D reconstruction of a E16.5 IFT140^null1/null1^ embryo processed using episcopic confocal microscopy highlighting abnormal chest/lung development. dAo: Descending aorta, L: Liver, SCV: Subclavian vein, SVC: Superior vena cava, TEF: Tracheoesophageal fistula, UL: Underdeveloped Lung.
